# Energy Constraints and Neural Strategy Transitions in Alzheimer’s: A Game-Theoretic Model

**DOI:** 10.1101/2025.05.24.655918

**Authors:** Irina Kareva, Georgy Karev

## Abstract

While many mechanisms have been proposed to drive Alzheimer’s disease, particularly the accumulation of amyloid plaques and hyperphosphorylation of tau proteins, emerging evidence suggests that they may be the byproducts of earlier damage rather than initiating events. Instead, metabolic dysfunction and the inability of neural cells to support their energetic demands may be a more plausible trigger for subsequent pathological cascade (the neuron energy crisis hypothesis). Here we highlight how type 2 diabetes (T2D) can contribute to neurodegeneration by impairing brain energy metabolism. We present a game-theoretic framework, where neurons face trade-offs between energy efficiency and information fidelity. We show that under metabolic stress, neural networks can evolve toward smaller group sizes that prioritize energy efficiency over information quality, which may underlie the observed collapse of cognitive capacity during neurodegeneration. We conclude with a discussion of interventions, ranging from antidiabetic drugs to cognitive engagement and sensory stimulation, aimed at reducing metabolic stress and preserving cognitive function.

## Introduction

Alzheimer’s disease (AD) is the most common form of dementia. Globally, over 50 million people are currently living with dementia, with this number expected to triple in the next 30 years as the world population ages (Javaid et al. 2021). The likelihood of developing AD increases dramatically with age, with nearly a third of individuals over 85 affected to some extent (Association 2024). Comorbidities such as type 2 diabetes, cardiovascular disease, and metabolic syndrome are also strongly associated with increased dementia risk (Raffaitin et al. 2009; Lanctôt et al. 2024).

Over the years, numerous hypotheses have emerged to explain what initiates and drives AD. The most dominant models have focused on amyloid-β accumulation and tau hyperphosphorylation, leading to plaques and tangles, respectively. However, both anti-amyloid and anti-tau therapies have yielded only modest clinical benefits, often slowing cognitive decline by mere months, if at all (Van Dyck et al. 2023; Wojtunik-Kulesza et al. 2023; Boxer et al. 2019; Teng et al. 2022). These outcomes, combined with emerging evidence that both amyloid-beta and tau may serve evolutionarily conserved protective roles, such as antimicrobial defense and cytoskeletal repair (Robert D Moir et al. 2018; Spillantini & Goedert 2013), suggest that their accumulation may be a downstream effect of earlier cellular or systemic dysfunction rather than the root cause.

In recent years, the metabolic hypothesis of Alzheimer’s has gained traction as an alternative to the traditional protein-centric models that emphasize amyloid-β and tau as primary causes. Instead, this hypothesis suggests that neuronal energy failure precedes and potentially initiates downstream pathological cascades. Positron emission tomography (PET) imaging studies have demonstrated that glucose hypometabolism in regions such as the posterior cingulate and temporoparietal cortex can be detected well before amyloid plaque formation (Mosconi et al. 2008), and mitochondrial dysfunction is consistently observed in AD-affected brains (D’Alessandro et al. 2025; Dewanjee et al. 2022; Butterfield & Halliwell 2019). In a particularly revealing experiment, Berndt et al. (Berndt et al. 2001) showed that partial inhibition of cytochrome c oxidase (Complex IV of the mitochondrial electron transport chain) with sodium azide produced memory impairments in animal models that mirrored those seen in Alzheimer’s, without any amyloid accumulation. This suggests that at least in some cases neurodegeneration can begin with energy failure, not protein toxicity.

These findings are in line with an increasingly common description of Alzheimer’s as “type 3 diabetes”, a condition marked by brain-specific insulin resistance and impaired glucose utilization (González et al. 2022; Monte & Wands 2008). This analogy highlights the deep metabolic underpinnings of AD, particularly the observation that individuals with Type 2 Diabetes (T2D) are at significantly higher risk for developing Alzheimer’s (Barbagallo & Dominguez 2014). Yet paradoxically, despite the hallmark systemic hyperglycemia of T2D, many of these individuals exhibit signs of glucose deprivation in the brain. This counterintuitive phenomenon is central to the model explored in this paper.

To understand this paradox, it is essential to first understand how the brain accesses glucose. Glucose enters the brain across the blood-brain barrier (BBB) via insulin-independent GLUT1 transporters (Simpson et al. 2007), which ensure that even under conditions of low systemic glucose, the brain receives a stable nutrient supply. However, chronic hyperglycemia, as seen in T2D, disrupts this balance. Long-term exposure to high glucose levels has been shown to impair both the function and localization of GLUT1 transporters (Duelli & Kuschinsky 2001), reducing the brain’s capacity to import glucose despite its systemic abundance.

Moreover, hyperglycemia damages the endothelial cells that make up the BBB, leading to its increased permeability, often referred to as a “leaky” BBB (Stanimirovic & Friedman 2012). This allows toxins and inflammatory cytokines to enter the brain, while simultaneously destabilizing the regulatory mechanisms that normally maintain steady glucose influx. As a result, glucose delivery to the brain becomes erratic, oscillating between brief periods of overload and prolonged periods of deprivation. The net effect is that, over time, neurons may receive less functional glucose than they would under normoglycemic conditions.

These disruptions are further exacerbated by hyperglycemia-induced formation of advanced glycation end products (AGEs), which provoke oxidative stress, inflammation, and the loss of small blood vessels in the brain, also known as capillary rarefaction (Brownlee 2001; Prakash & Carmichael 2015). The result is a decline in cerebral perfusion, meaning that even when glucose is technically present in the extracellular fluid of the brain, it may not be adequately delivered to neurons. This decoupling between systemic glucose availability and neuronal energy supply helps explain why T2D increases the risk for cognitive decline despite apparent energy abundance.

The consequences of this mismatch are especially severe for neurons, which are highly sensitive to energy instability. Unlike liver or muscle cells, neurons cannot store glucose or rely on glycolysis; instead, they are almost entirely dependent on oxidative phosphorylation via the TCA cycle, requiring a continuous, finely regulated glucose supply. When this supply becomes unreliable, it can lead to mitochondrial failure.

Importantly, this energy instability can also initiate a self-perpetuating feedback loop of damage. Mitochondrial dysfunction leads to the release of mitochondrial damage-associated molecular patterns (mtDAMPs), which activate microglia and trigger inflammatory responses that further damage neuronal networks (Freeman & Ting 2016; Heneka et al. 2018). As the system spirals, amyloid-β and tau, which may initially play protective roles by sealing BBB leaks or restructuring cytoskeletons under stress (Robert D. Moir et al. 2018; Iqbal et al. 2010), begin to accumulate pathologically, further impairing mitochondrial dynamics and synaptic function. A summary of the key processes underlying the cascade of damage triggered neural energy crisis model are given in Figure 1.

**Figure 1.**
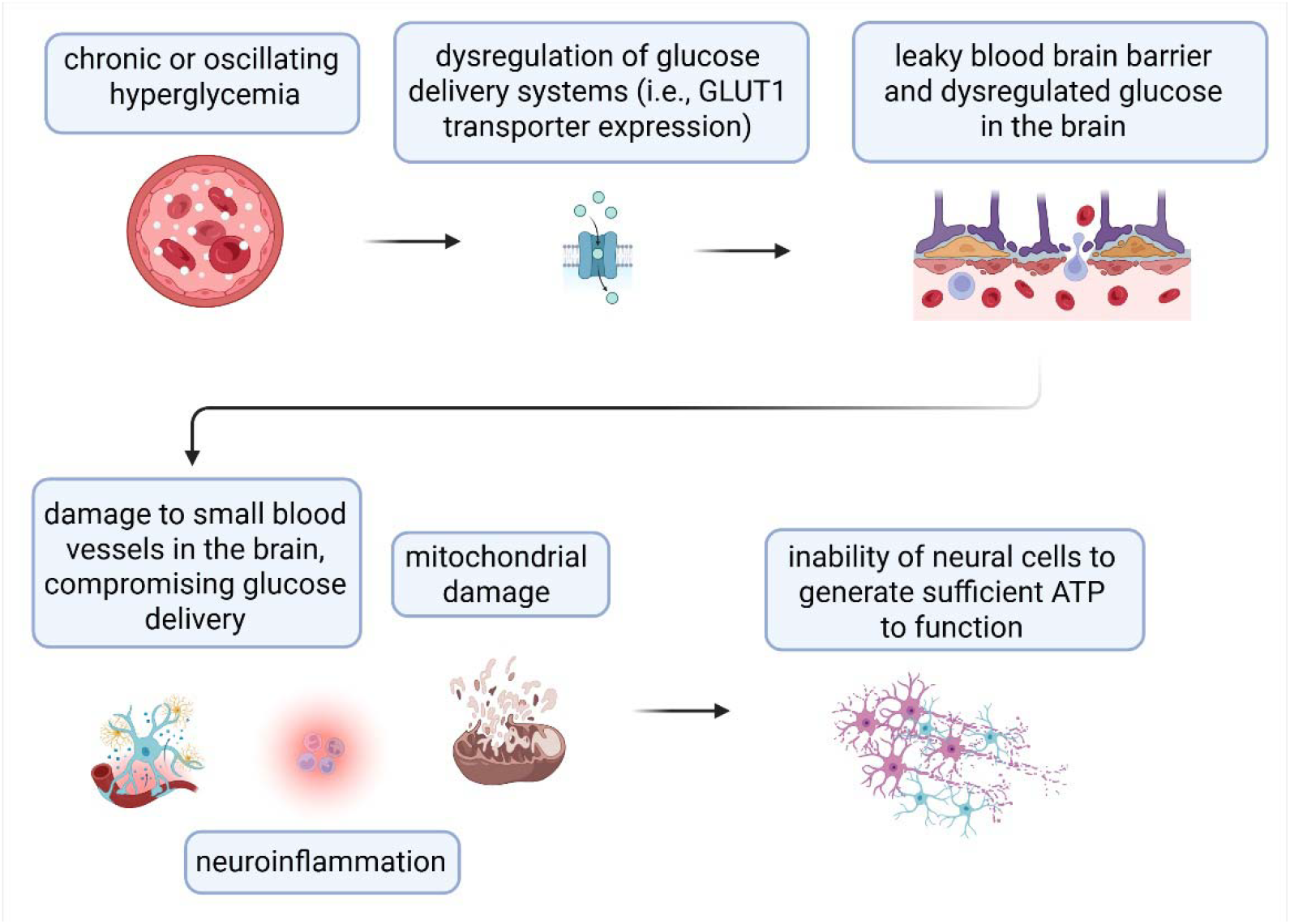
Key mechanisms underlying hyperglycemia-induced neurodegenerative cascade.

Different mathematical modeling approaches have been employed to capture the emergence and progression of cognitive decline and dementia. Multidimensional ODE models have been used to simulate the temporal evolution of biomarkers and cognitive tests, allowing for personalized prognostic predictions by capturing disease progression pathways in Alzheimer’s disease (Bossa & Sahli 2023). Biophysical factor models have also been applied to explore the role of various interconnected biological processes in the onset and evolution of AD. These models incorporate factors such as amyloid-beta aggregation, tau hyperphosphorylation, and neuronal loss (Moravveji et al. 2024). Other ODE-based approaches have focused on modeling neuroinflammation and protein dynamics by capturing interactions between neurons, astrocytes, microglia, and peripheral macrophages (Hao & Friedman 2016). These models have predicted the effects of potential therapeutic interventions, suggesting that targeting neuroinflammatory pathways may mitigate disease progression. Some other approaches to this topic include looking at it through the lens of game theory. For instance, evolutionary game theory on networks (EGN) has been used to model brain network dynamics, providing insights into the stability and adaptability of neural interactions in the presence of disease-related disruptions (Madeo et al. 2017).

Here we explore the consequences of metabolic disruptions through the lens of game theory and group size selection subject to metabolic constraints. In states of energetic abundance, neurons may participate in large, information-rich networks that support complex cognitive functions. However, when energy becomes scarce or unstable, neurons may retreat into smaller, low-cost subgroups, preserving survival at the expense of information fidelity. In our analysis, we explore ways to influence these transitions as a function of emerging hyperglycemia and T2D. We conclude with a discussion of available intervention methods that may delay this transition in an effort to preserve cognitive function.

## Model description

### Let us consider the following framework

Assume that during neural interactions, there exists a trade-off between energy efficiency E and information transmission (I). There exist energetic constraints on neural firing, and trade-offs between maintaining a high signal-to-noise ratio (SNR) bandwidth, and the energy costs of neural activity (Laughlin 2001). That is, assume that a neuron’s firing strategy is inherently shaped by the tension between conserving metabolic energy and ensuring efficient and reliable information transmission.

Group size plays a key role in this tradeoff. Larger groups allow more robust signal representation but are metabolically costly; smaller groups conserve energy but at the expense of signal fidelity (Laughlin 2001). One of the questions that we would like to explore here is whether in ample resources, we’d observe selection for larger groups, and if, as the energy supply to neurons becomes scarcer, we’d observe selection for smaller groups.

In order to achieve this goal, we will build on the work by (Karev 2024), where the author studies which group size will be selected over time as a function of the payoff matrix that governs interactions between two individuals playing two possible strategies. The author tracks the evolution of the distribution of parameter *β*, which is defined as being inversely proportional to the fixed size *N* of subset of the population, with which each individual can interact. That is, if 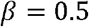, then an individual has available for interaction of only one another individual. If 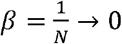, then the number of possible interaction partners is indefinite. Notably, the exact composition of this subset N of the entire population that is available for interaction is random and can be composed of individuals using either strategy or any combination thereof. The outcome of the interaction is determined by the values of the coefficients of payoff matrices. Parameter *β* here refers only to the size of the subpopulation, from which an interaction partner can be selected. A summary of the key calculations necessary to implement this approach is provided in the Appendix A.

Within the frameworks of this work, a sample payoff matrix for the interactions between neurons would have the following form:

|  | Neuron B: Energy Efficiency | Neuron B: Information transmission |
| --- | --- | --- |
| Neuron A: Energy Efficiency | a | b |
| Neuron A: Information transmission | c | d |

Since both energy efficiency and information quality are important for neural function, we introduce the concept of a relative benefit, or *utility* derived from each strategy, where a more energy efficient strategy results in lower information quality, and a higher information quality strategy also carries energetic costs. Here the utility of each strategy is relative and is derived from balancing trade-offs, rather than being an absolute measure of either energy cost or information quality.

The first basic formulation of a utility function could be as follows:

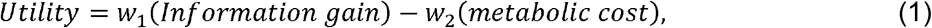

where *w*_1_ and *w*_2_ are weights that would reflect the priority of one vs the other in any given context. A calculation of sample coefficient values is given in Appendix B.

Within the framework of our model, neuron group size described by parameter β would vary between 0 (large group size) and 0.5 (small group size). For our model, therefore, we’d expect the baseline case to tend to β →0, where each neuron has a large number of other neurons that are available for interaction (see Figure S1, Case 0).

Next, we would like to evaluate, whether it is possible to affect system dynamics in such a way as redirect population evolution from large groups sizes to small? We hypothesized that this can be achieved by changing the signs in the relative elements of the payoff matrix, similarly how changing the relative values in the top element of payoff matrix in the classical two-player Hawk-Dove game can change the outcome of the interaction away from mixed strategy to pure strategies, effectively turning the game into a Prisoner’s Dilemma. The results of this analysis are shown in Appendix C, confirming that indeed, selectively changing the sign of one or two elements in the payoff matrix can redirect group size evolution (Figure S1).

### Introducing the Switcher: Gradually Changing the Sign of Elements in the Payoff Matrix

Now that we have demonstrated that a simple sign change can lead to change in evolution of group size from large to small, we will introduce a “switcher function”, where such a change can occur more smoothly. For this purpose, we propose the following functional form:

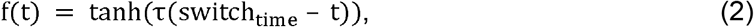

where parameter *τ* affects the steepness of the curve, and parameter switch_time_ affects the point in time, at which the sign of the coefficient switches from a positive value to a negative one. An illustration of the dynamics of the switcher function over time, with *τ* = 0.1 and switch_time_= 100, is shown in Figure 2A.

**Figure 2.**
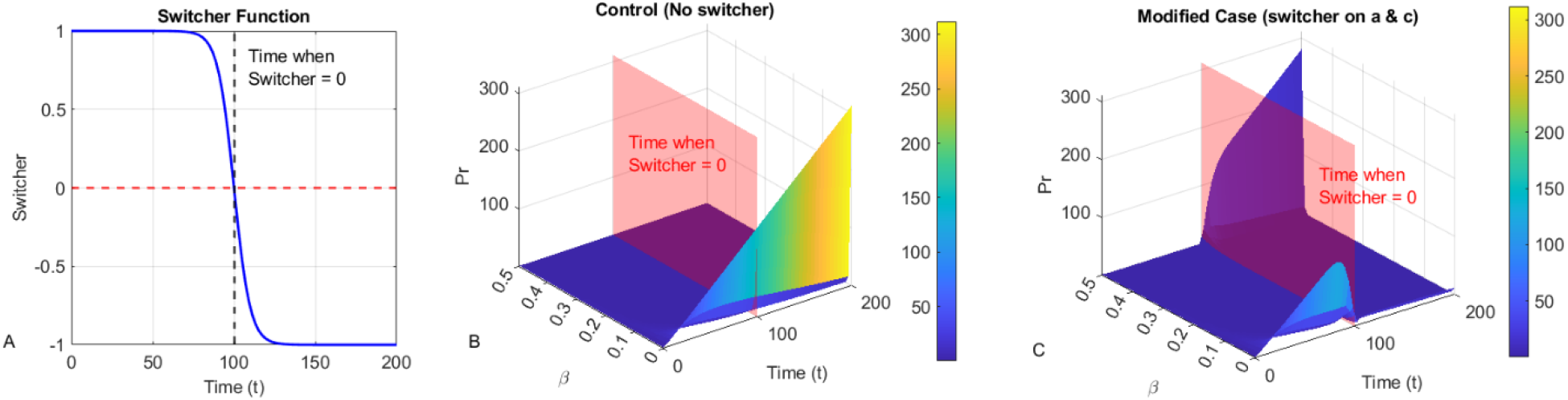
The impact of the switcher function. (A) The dynamics of the switcher f(t) over time, showing how it crosses zero at t=100. (B) Dynamics of distribution of the parameter *β* in the control case, when the switcher is not coupled with any coefficients of the payoff matrix [a b; c d] but the time that it crosses zero is still shown to highlight the difference the next case. (C) Dynamics of distribution of the parameter *β* in the modified case, when the switcher is placed on the coefficients a and c of the payoff matrix [*af*(*t*) *b*; *cf*(*t*) *d*]; the time when the switcher crosses zero (here at *t* = 100) is shown with the transparent plane, highlighting the difference in trajectories compared to the control case.

Such a function can be coupled to the coefficients of the payoff matrix, resulting in a gradual transition from positive to negative coefficient values. An example of the modified payoff matrix, where the switcher is placed on coefficients a and c, would be as follows:

|  |  |  |
| --- | --- | --- |
|  | **Neuron B: Energy Efficiency** | **Neuron B: Information transmission** |
| **Neuron A: Energy Efficiency** | $af(t)$ | $b$ |
| **Neuron A: Information transmission** | $cf(t)$ | $d$ |

As one can see in Figure 2C the switcher function starts decreasing around time *t=75*, reducing the values in the payoff matrix of the *af*(*t*) and *cf*(*t*). Consequently, the distribution of the parameter *β* also starts to change even prior to the abrupt sign shift, signaling the upcoming transition, in contrast to the case with no switcher (Figure 2B). After the switcher crosses zero, here at *t=100*, the distribution of group sizes continues much more rapidly.

### Modifying the Switcher to be a Function of Fasting Glucose Levels

Now, we are ready for the main step, which is showing how hyperglycemia associated with type 2 diabetes could result in such a switch, causing transition of the neural group size from large (β→0, predicted to be prioritizing quality of information over energy costs) to small (β→0.5, predicting to prioritize energy costs, even at the expense of quality of information transmission).

Recall that there paradoxically exists a positive correlation between high systemic glucose levels and low glucose availability for brain cells: as baseline glucose increases over time as an individual may be developing pre-diabetes and diabetes, at some concentration it may become damaging enough to trigger the pathological cascade that eventually leads to neural energy crisis (high systemic glucose → formation of AGEs and dysregulation of glucose delivery systems and GLUT1 transporter expression → leaky BBB → damage to small blood vessels to compromise glucose delivery to cells + damage to astrocytes as sources of additional energy + mitochondrial stress → inability of neural cells to generate sufficient ATP to function). While we will not model all the steps of the pathological cascade and feedback loops, we can capture the key aspects of the process as follows: increase in glucose → surpassing some threshold that triggers the pathological cascade → increased metabolic costs for neural cell function (since low availability of functional glucose or inability to utilize it effectively translates into a state of nutrient deprivation).

Within the frameworks of our model, we can therefore modify the switcher function in the following way:

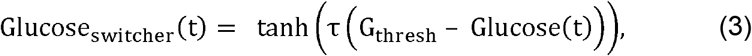

where the sign switch will occur when Glucose(t) = G_thresh_, such that when baseline concentrations of glucose exceed the G_thresh_ value, the switcher function changes signs. The payoff matrix then becomes as follows:

|  | **Neuron B:**<br>**Energy Efficiency** | **Neuron B:**<br>**Information transmission** |
| --- | --- | --- |
| **Neuron A:**<br>**Energy Efficiency** | $a * \text{Glucose}_{\text{switcher}}(t)$ | $b$ |
| **Neuron A: Information**<br>**transmission** | $c * \text{Glucose}_{\text{switcher}}(t)$ | $d$ |

Next, in order to evaluate the impact of the glucose-dependent switcher on neural group size dynamics, we first need to describe how glucose concentrations change over time, and how emergence of type 2 diabetes can lead to this switch. For this purpose, we will adapt a mathematical model developed in (Kareva 2024).

In summary, this model captures emergence of insulin resistance and hyperglycemia as a function of decreasing capacity to store glucose, described by variable K(t). As the capacity to store glucose decreases due to, for instance, age-related muscle loss, baseline concentrations of extracellular glucose start to increase, capturing slow emergence of pre-diabetes and diabetes over time. When fasting glucose levels cross the threshold G_thresh_ (which can be any representative level of glucose, i.e., 7.5 mM; the exact value is likely individual-specific), the switcher function will cross zero, and the coefficient in the payoff matrix will change signs.

A description of the equations and underlying assumptions of the glucose-insulin model, as well as parameter Table and a representative simulation of underlying glucose-insulin dynamics, can be found in the Appendix D and in (Kareva 2024).

### Now let us put these pieces together

We now have a dynamical system that is capturing gradual increase of baseline systemic glucose levels over time, described by dynamic variable Glucose(t). The value of Glucose(t) at each time point is coupled with the proposed switcher function as described above. The switcher is coupled with the utility payoff matrix as described above. We are now ready to analyze our final system and ask the following question: can we capture the transition from large neuron group sizes to small ones as a function of increasing systemic hyperglycemia?

For that, we start with the coefficients calculated for a utility payoff matrix as described above. If a switcher, which is restricted to [−1 1], is placed on a coefficient, it becomes

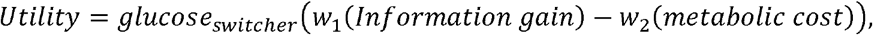

The baseline case has the utility coefficient be positive (switcher = 1); however, as glucose concentration increases to G_thresh_, switcher function crosses zero and becomes negative, changing the sign of the utility coefficient as a function of glucose concentration.

Let us now show the dynamics of such a system, were the glucose-dependent switcher on elements a and c (notably, it can be placed on other coefficients as well, and this pair here was chosen for illustration).

### The result of these simulations is shown in Figure 3

In the simulations done in Figure 3 we track the following metrics: the heatmap represents the change in the distribution of parameter β over time (recall that *β*= 1/*N*, where N is the total population size; β → 0 indicates evolution towards a large group size, while β → 0.5 indicates evolution towards a small group size). We also keep track of change in glucose carrying capacity K, fasting glucose, fasting insulin, and the value of the switcher function, as well as the time at which the switcher crosses the zero threshold and flips the sign of the coefficient in the payoff matrix.

**Figure 3.**
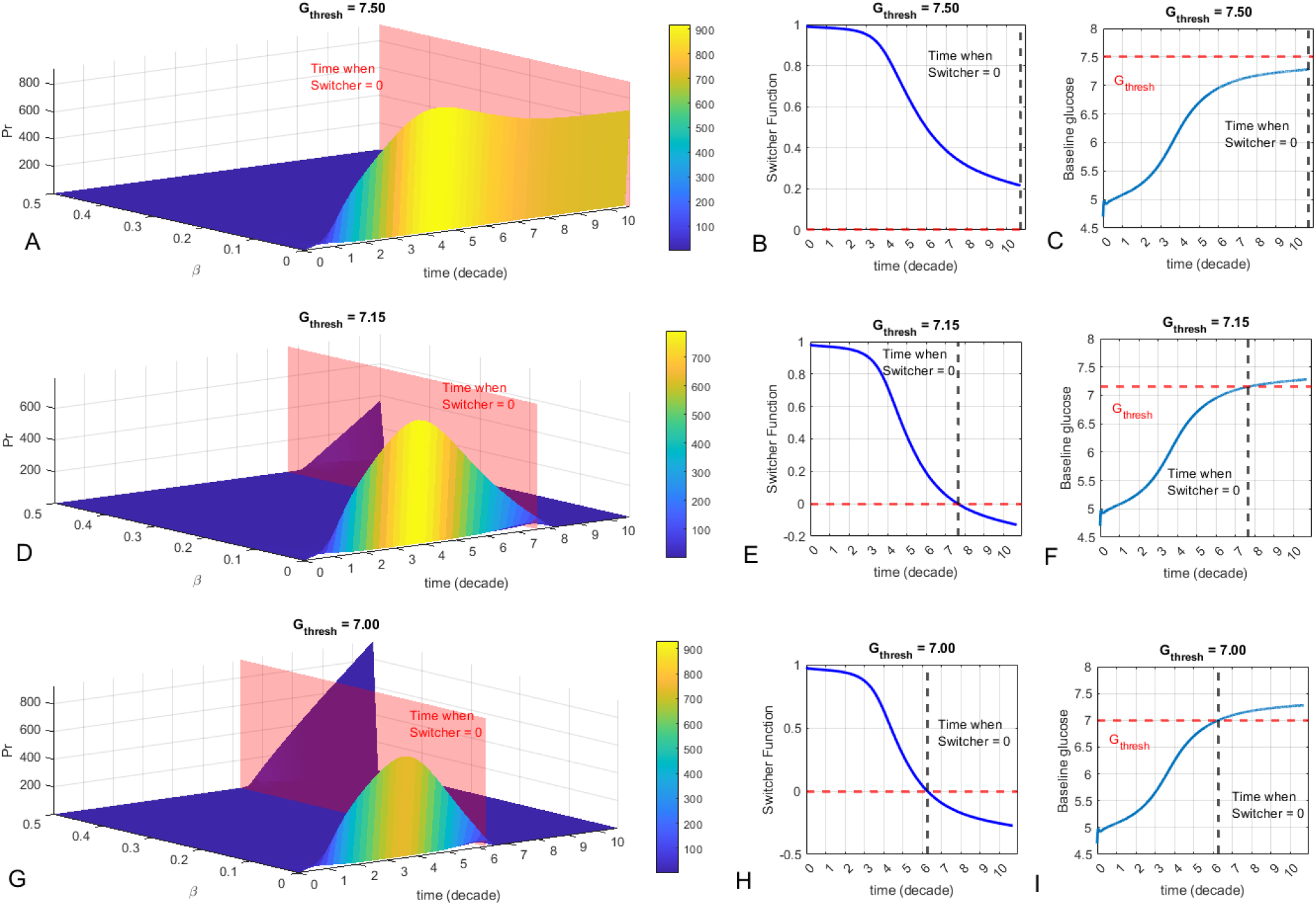
The impact of varying glucose-dependent thresholds in the switcher function on evolution of group size over time. (A) G_thresh_ = 7.5 mM. The change over time of the distribution of parameter β. (B) The dynamics over time of the glucose switcher function, which never reaches zero. (C) The dynamics over time of glucose, which never reaches the threshold of 7.5 mM. (D) G_thresh_ = 7.15 mM. The change over time of the distribution of parameter β; the transition from larger to smaller groups happens around 75 years old. (E) The dynamics over time of the glucose switcher function, which crosses zero at approx. 75 years old. (F) The dynamics over time of glucose, which reaches the threshold of 7.15 mM at approx. 75 years old. (G) G_thresh_ = 7.0 mM. The change over time of the distribution of parameter β; the transition from larger to smaller groups happens around 60 years old. (H) The dynamics over time of the glucose switcher function, which crosses zero at approx. 60 years old. (I) The dynamics over time of glucose, which reaches the threshold of 7.0 mM at approx. 60 years old. Utility matrix payoff coefficients used in this simulation are a = 0.6, b = 0.03, c = 0.7, d = 0.02. The baseline values of the coefficients were selected for illustration purposes and can be adjusted based on context specific considerations, i.e., as described in Appendix B.

In the top panel of Figure 3 we show the case, when the “switch” level is never reached. The value of the coefficient may be decreasing over time, since the switcher function is decreasing, but it never changes sign, and the distribution of strategies with respect to group size does not undergo any abrupt changes.

In the middle panel, we set the glucose switch threshold at 7.15 mM. One can see that in this model this occurs around 75 years old, at which point the switcher function crosses zero. We can now observe a qualitative change in dynamics of distribution of *β* at this time. Finally, if one’s threshold occurs when glucose is 7 mM, the switch occurs even earlier, in this model at around 60 years old. (While the values of parameters used in these simulations are representative and not quantified based on data, it is interesting to observe that the “peak” number of neural connections are predicted to occur around 20-40 years old, with subsequent declines occurring slower or faster depending on the rate of decline of the switcher function).

Interestingly, the model predicts the “slow decline” period before the critical transition in the switcher function; however, after the switcher crosses zero, the change then occurs very quickly. This is consistent with clinical observations of slower prodromal stages that may take years (Caldwell et al. 2015; Association 2024), followed by rapid decline when a critical threshold is passed.

## Discussion

In this paper we propose a game-theoretic framework for understanding cognitive decline as an adaptive, emergent property of metabolic stress in the brain. Within this model, neurons interact under constraints of limited energy and competing demands for information fidelity. The results show that, under increasing systemic glucose concentrations, such as those seen in type 2 diabetes (T2D), neurons may shift from participating in large, high-fidelity networks to forming smaller, energy-conserving groups, potentially resulting in behavioral phenotypes consistent with cognitive impairment.

The model introduces a switcher function, a smooth threshold-based transition that alters the utility payoff of different neural strategies. When baseline glucose exceeds a (likely individual-specific) threshold, this function causes a shift in the relative value of information-rich versus energy-efficient behavior. Simulations demonstrate that this can trigger a phase transition in neural group dynamics, with expected group sizes declining sharply after surpassing the threshold. Importantly, this mirrors clinical observations in Alzheimer’s disease, which often involves a prolonged prodromal phase followed by accelerated cognitive decline (2, 39).

While the current implementation uses a glucose-based switcher for proof of concept, the framework is flexible. In future iterations, multiple switchers could be introduced to reflect diverse biological, environmental, or behavioral stressors. For example, one switcher might reflect hyperglycemia and impaired glucose transport, while another could represent neuroinflammation, oxidative stress, or traumatic injury. Moreover, different elements of the payoff matrix could be modulated independently to reflect the heterogeneous effects of these stressors on different neuronal populations or between different individuals.

This adaptability provides flexibility for exploring how therapeutic interventions might influence system dynamics. For instance, interventions that improve energy metabolism, such as SGLT2 inhibitors or GLP-1 receptor agonists, have already shown potential to reduce dementia risk in T2D populations (Shin et al. 2024; Nørgaard et al. 2022; Tang et al. 2023). These therapies might be modeled directly as delaying the switcher’s crossing of the threshold or incorporated into the underlying glucose-insulin model that governs glucose dynamics over time, effectively extending the period during which neurons can sustain larger group sizes and higher information fidelity.

Similarly, interventions that increase cognitive load or sensory stimulation, such as cognitive training, hearing aids, or vision correction, could be modeled as increasing the relative weight of information quality in the utility function. This would shift neural strategy selection toward larger groups, reinforcing synaptic connections and possibly counteracting energy-conserving behaviors. Empirical studies support this: sensory deficits are known to increase cognitive load and accelerate decline, while correcting them appears to reduce this risk (Cantuaria et al. 2024; Lin et al. 2023).

Other emerging therapies directly target the mitochondrial dimension of the neuronal energy crisis. For example, photobiomodulation (PBM), which uses near-infrared light to stimulate cytochrome c oxidase (Complex IV), enhances ATP production and reduces oxidative stress (Wang et al. 2016; Nairuz & Lee 2024). Intranasal insulin has also shown promise in improving cognitive performance in patients with mild cognitive impairment, presumably through supporting the function of astrocytes, which in turn support neural function (Fernandez et al. 2022; Craft et al. 2012). These could be modeled as interventions that diminish the metabolic cost term, allowing for higher fidelity signaling without the same energetic burden.

The model can also account for how pathological processes like infection, trauma, or inflammation increase energy costs and drive network collapse. For instance, chronic infections such as HSV-1 (Linard et al. 2020), Lyme disease (Parthasarathy & Gadila 2022), or Porphyromonas gingivalis in periodontal disease (Dominy et al. 2019) have all been linked to increased dementia risk via inflammatory and mitochondrial pathways. Severe systemic infections (Bohn et al. 2023) or post-viral syndromes like long COVID (Talkington et al. 2025) may produce similar effects by increasing neuroinflammation and disrupting the blood-brain barrier. All of these mechanisms could be formalized in future models as factors increasing the slope or magnitude of the switcher function, accelerating cognitive decline in susceptible individuals.

One of the key strengths of this modeling framework is its potential to inform personalized medicine. By formalizing how biological stressors and therapeutic interventions influence neural group dynamics, this model can be used to simulate individual trajectories based on biomarker data. For example, an individual with insulin resistance and hearing loss might be modeled with a faster-acting switcher and lower information weights, while another with high cognitive reserve and good cardiovascular fitness might show greater resilience to metabolic stress.

Finally, the model’s simplicity leaves ample room for expansion. Switchers can be made time-dependent, dynamically responsive to interventions, or specific to neuron subtypes or brain regions. Future models could incorporate feedback loops, disease progression pathways, or even stochastic perturbations that reflect real-world biological variability. With the rise of wearable technologies capable of detecting early motor changes or cognitive shifts (Obuchi et al. 2024; Zhang et al. 2024), such models may soon support early-warning systems for identifying individuals at risk of transitioning from adaptive compensation to irreversible decline.

In sum, this work lays the foundation for a systems-level view of Alzheimer’s disease – one that reframes cognitive decline as a shift in neural strategy rather than a consequence of protein accumulation. By tying neural decision-making to energetic constraints, this framework opens new avenues for understanding, predicting, and potentially delaying the onset of dementia, leveraging both existing and emerging interventions.

## Appendix A. Summary of the background theory for tracking evolution of groups sizes by G. Karev (2024).

Let us describe the initial model of two-player games with two possible strategies, that we denote as 0 and 1. Denote *x*_*i*_ (*t*) the frequency of *i*-th strategy, *i*=0,1. Let 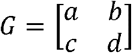 be the payoff matrix of a game. Then the dynamics of this game is given by the replicator equation (see, e.g., (Hofbauer & Sigmund 1998; Taylor & Jonker 1978)):

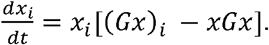

Consider a family of games with corresponding payoff matrices *G*^*β*^ parametrized by the parameter *β*. Consider a model of a community composed of populations with different payoff matrices *G*^*β*^ and study the natural selection between the populations. This process can be formally described by the evolution of the distribution of the parameter *β*.

Classical replicator dynamics assumes that each individual interacts with a representative sample of the infinite population. In (Hilbe 2011), the author considered the *local replicator dynamics*: while the population itself is infinite, interactions and reproduction occur in random groups of size *N*.

Hilbe proved that if groups are formed according to a multinomial distribution, then the local replicator dynamics is given by the *modified replicator equation*

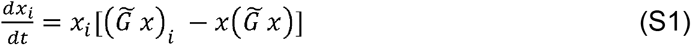

With 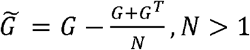.

Let us denote 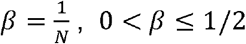 and represent the matrix 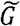 in the form

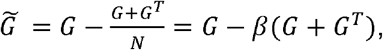

Denote

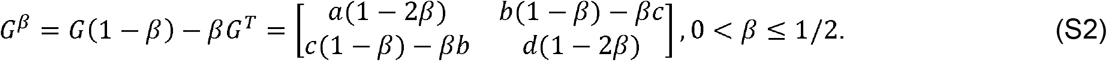

Then, according to (Hilbe 2011), the local replicator dynamics at given value of 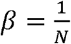 is described by the replicator equation

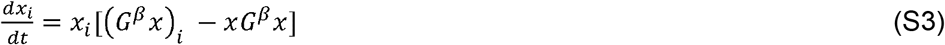

with the parametrized payoff matrix given by Equation (S2).

The question we are interested is the dynamics of distribution *P*(*t,β*) of the parameter *β*. To answer this question we use the following results given in (Karev 2024).

Assume that strategies 0 and 1 are chosen at the initial time with corresponding probabilities *p*_0_ and p_1_ =1 − *p*_0_ for all values of *β*. The initial distribution *P*(0,*β*) of parameter *β* is taken as exponential truncated on the interval (0,1/2):

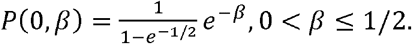

Define the variable *q*(*t*) by the equation

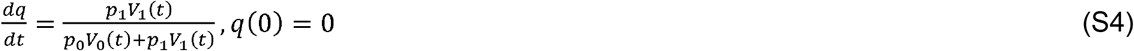

Where

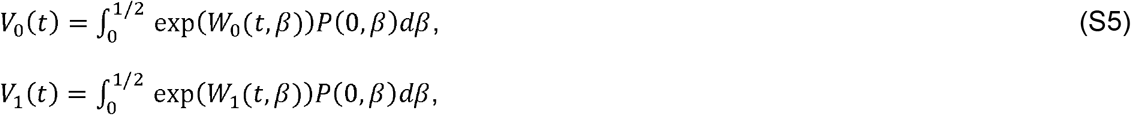

and

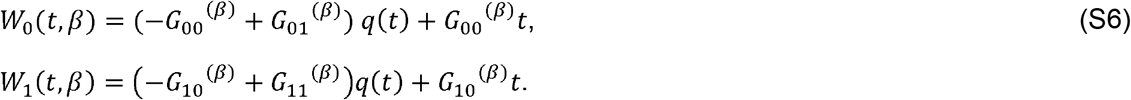

Then

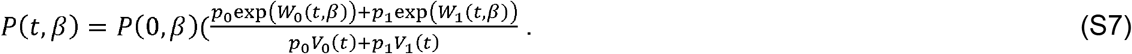

Equations (S4-S7) allow us to compute the distribution of the parameter *β* for all *t* and hence to trace the dynamics of group sizes *N*.

## Appendix B. Coefficient Derivation and Normalization

Let us explore how we could estimate coefficients to populate our utility payoff matrix. We know that actively firing neurons are expected to have high energetic demands for spiking and synaptic transmission. Resting neurons, on the other hand, are expected to have much lower metabolic demands. With respect to information gain, we’d expect it to be minimal during resting state, and significant for an active state. Restricting these coefficients on the scale from 0 to 10, we can expect the following baseline coefficients: Energy Cost is 1 for the Energy Efficient strategy and 10 for the High Information Strategy; Information Gain is 0 for the Energy Efficient strategy and 10 for the High Information Strategy.

Putting this information together, let us now formulate a payoff matrix that we will take forward. Recall that the “payoff” considered here is taken as utility, defined as follows:

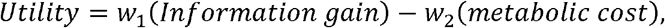

where *w*_1_ is the weight for information gain, emphasizing its importance in the context, *w*_2_ is the weight for energy cost, reflecting metabolic constraints, Information Gain is a measure of signal fidelity, throughput or SNR, and Energy Cost is the metabolic expenditure in ATP. Contextually, we expect *w*_1_>*w*_2_ in high-demand contexts, where information fidelity is prioritized, while we expect *w*_2_>*w*_1_ if contexts that prioritize energy efficiency.

For a high demand context, sample values of the weights could be *w*_1_ = 0.7, and *w*_2_ = 0.3; these values reflect critical importance of fast, reliable information transmission, with energy costs perhaps being less critical in the short term. The payoff matrix then becomes:

| High energy context,<br>Raw utility values | Neuron B: Energy<br>Efficiency | Neuron B: Information<br>transmission |
| --- | --- | --- |
| Neuron A: Energy<br>Efficiency | $0.7*0 - 0.3*1 = -0.3$ | $0.7*10 - 0.3*1 = 6.7$ |
| Neuron A: Information<br>transmission | $0.7*0 - 0.3*10 = -3$ | $0.7*10 - 0.3*10 = 4$ |

For convenience, we normalize these coefficients to be in the range between [0 1] (the method we use is described below), we get the following matrix for the high energy context:

| High energy context,<br>Normalized to [0 1] | Neuron B: Energy<br>Efficiency | Neuron B: Information<br>transmission |
| --- | --- | --- |
| Neuron A: Energy | 0.28 | 1 |
| **Efficiency** |  |  |
| **Neuron A: Information transmission** | 0.01 | 0.72 |

Similarly, for a low-energy context, we can take the coefficients to be *w*_1_ = 0.3 and *w*_2_ = 0.7, reflecting the importance of conserving energy over information fidelity (i.e., in an energy limited environment), resulting in the following payoff matrix:

|  |  |  |
| --- | --- | --- |
| **Low energy context, Raw utility values** | **Neuron B: Energy Efficiency** | **Neuron B: Information transmission** |
| **Neuron A: Energy Efficiency** | $0.3*0 - 0.7*1 = -0.7$ | $0.3*10 - 0.7*1 = 2.3$ |
| **Neuron A: Information transmission** | $0.3*0 - 0.7*10 = -7$ | $0.3*10 - 0.7*10 = -4$ |

Normalizing these coefficients to be between [0 10], we get the following matrix for the high energy context (the details of this calculation are given in the appendix):

|  |  |  |
| --- | --- | --- |
| **Low energy context, Normalized to [0, 1]** | **Neuron B: Energy Efficiency** | **Neuron B: Information transmission** |
| **Neuron A: Energy Efficiency** | 0.68 | 1 |
| **Neuron A: Information transmission** | 0.01 | 0.32 |

To normalize to a 0-1 scale, first let us take the range of raw utility values:

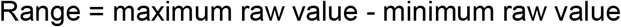

For the high demand context, minimum value is −3 and maximum is 6.7, resulting in a range of 9.7. For the energy limited context, minimum value is −7, while the maximum value is 2.3, resulting in the range of 9.3

Next, to normalize each raw utility value *x* to the desired range [a b], we can use the following formula:

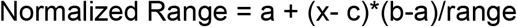

Where a is the minimum of the target range, b is the maximum, c is the minimum raw value (here −3 for high energy context and −7 for low energy context), and range is defined and calculated as above.

## Appendix C.

### Test Case

#### Changing the Sign of Elements in the Payoff Matrix

As a proof of principle exercise, let us take the matrix that was derived above and change the sign of the coefficients to see if the group size will change from large (β →0) to small (β →0.5).

That is, let this be the original matrix, where the values of the coefficients are normalized between 0 and 1 as described in the Appendix:

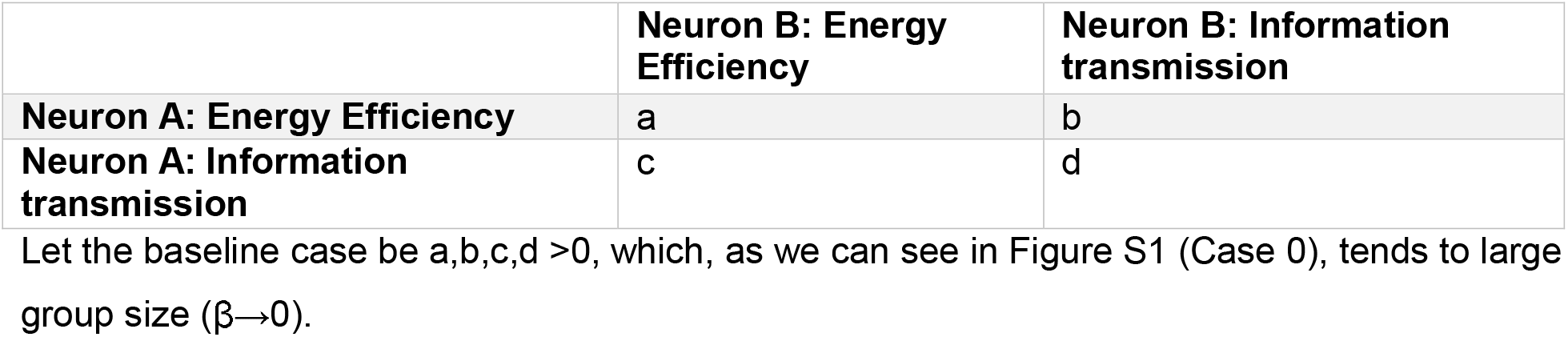

Let the baseline case be a,b,c,d >0, which, as we can see in Figure S1 (Case 0), tends to large group size (β→0).

Next, let us assess what happens with how β changes over time if we systematically change the signs on either one or two of the coefficients. For instance, in Case 1, we place a change of sign on coefficient a, so that a<0, b,c,d>0; in Case 2 we place the change of sign on coefficient b, such that b<0, a,c,d>0, etc. The full results of this numerical exercise are shown in Figure S1.

**Figure S1.**
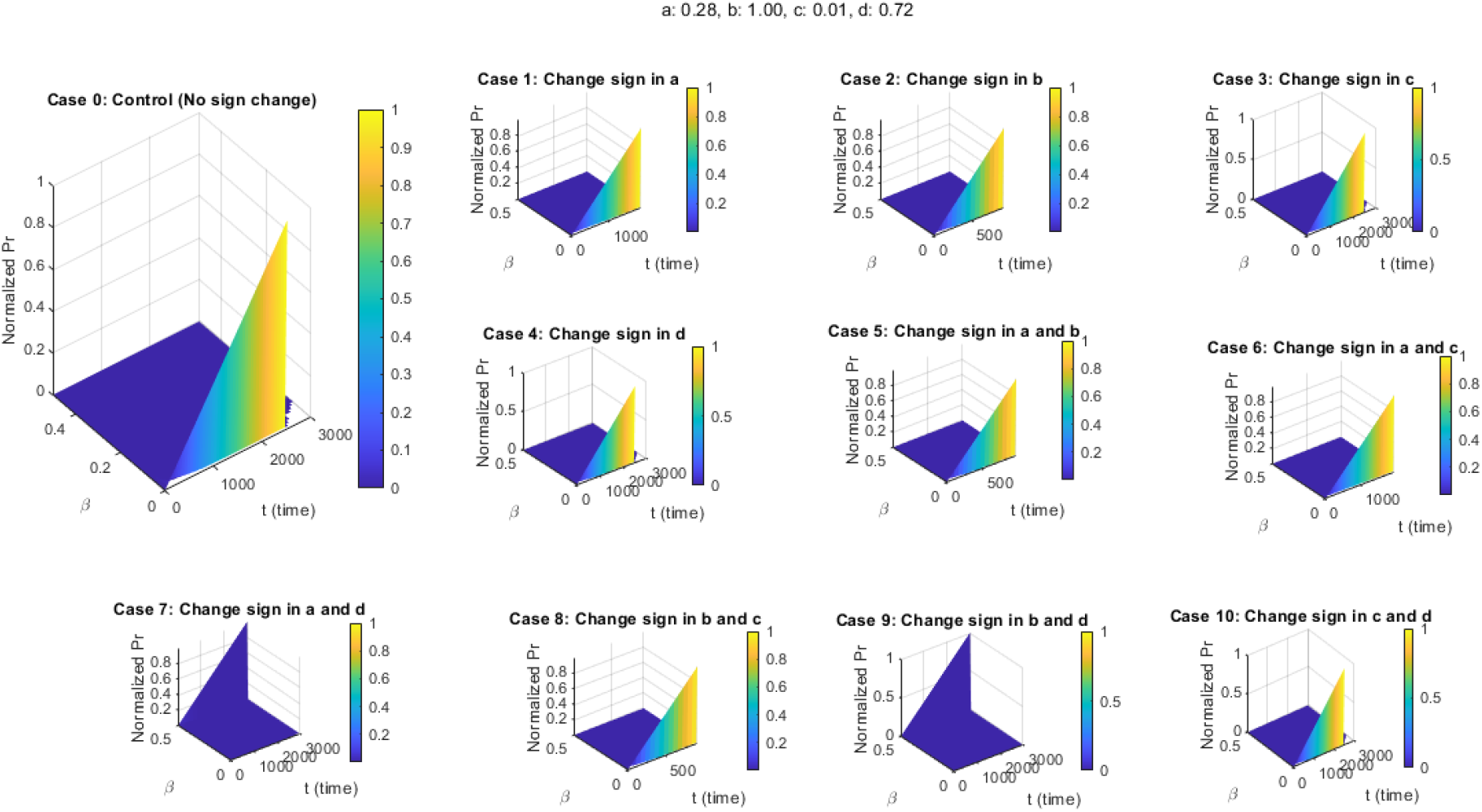
The impact of changing the sign of the coefficient of the payoff matrix on the evolution of group size over time. Depending on where the negative sign is placed, group size may remain as in the control case 0 (β →0, large group size), or change to β →1/2.

As one can see, for the payoff matrix with parameter values a = 0.28, b = 1; c = 0.01, and d = 0.72, there do exist several cases, when changing the sign can lead to change in the selected group size over time. Here, relative to control (Case 0), where all coefficients are positive, Case 7 (change in sign for a and d) and Case 9 (change in sign for b and d) result in qualitative changes in the overall selected group size, going over time from large groups (where we expect high importance to be placed on fidelity of information transmission) to smaller ones (where we expect to see energetic constraints, which would lead to lower quality of information processing).

Next, we wanted to evaluate whether these patterns are specific to these parameter values, or whether there exist more cases when placing a sign switch on the coefficients of the payoff matrix may lead to changes in group size in other regions of parameter space. For that we conducted a parameter sweep on all the coefficients (bound on the interval 0 to 1). We then calculated the following difference at the last time point tmax:

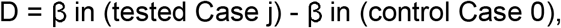

where j = 1…10.

If D = 0, then there was no difference in final group size (i.e., in both cases, β →0). If D > 0, then β in the test Case > β in the control case, and group size that is selected over time is decreasing (the larger the value of β, the smaller the group size). If D = 0.5, then the group has decreased to the smallest possible size.

#### The value of D for all the coefficients and all the test cases is reported in Figure S2

**Figure S2.**
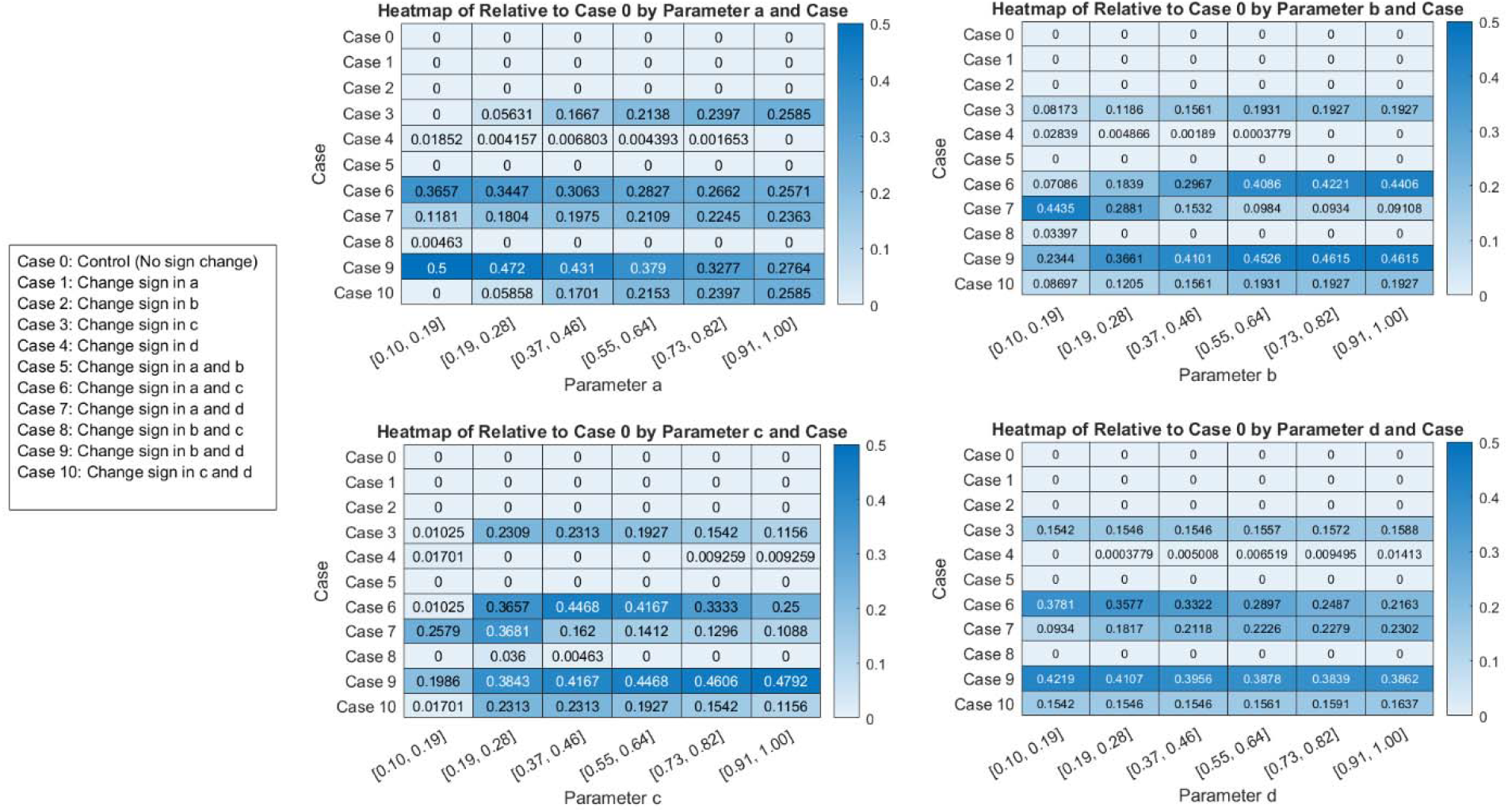
The impact of systematically changing the values of the coefficients in the payoff matrix on the difference between final group size relative to control for all the tested cases. Small difference (the value in the table) indicates that there is difference between the test case and the control case; large difference (up to 0.5) indicates that the average group size has evolved away from large and towards small groups.

As one can see in Figure S2, depending on the specific value of each parameter in the payoff matrix, it’s possible to change the final mean group size in some cases but not others. For instance, changing the sign of parameters a and b alone does not make a difference relative to control (Cases 1 and 2), as well as if we change the signs on both a and b (Case 5). Minor changes occur in cases 4 and 8 (change sign in d for Case 5, and in b & c in Case 8).

However, final group size can change dramatically, if we change the sign of coefficient c (Case 3), or in pairs of a & c (Case 6), a & d (Case 7), b & d (Case 9) and c & d (Case 10). Cases 6 and 9 (changing signs on coefficients a & c for Case 6, and b & d for Case 9) appear to have a particularly strong effect on the final group size.

## Appendix D. Key assumption of the glucose-insulin model by I. Kareva (2024)

The key mechanisms of the model introduced in (Kareva 2024) can be summarized as follows:

1) Blood glucose can increase from liver gluconeogenesis, which is inversely regulated by insulin (Hatting et al. 2018).
2) Glucose can be cleared in insulin-dependent and independent manner (i.e., via GLUT1, which is insulin-independent, or GLUT4, which is insulin dependent).
3) Insulin-dependent clearance involves binding between the insulin molecule and its receptor, which causes the translocation of the GLUT4 transporter on the surface of the cell, allowing the glucose to enter; rate of insulin-mediated glucose clearance is described by parameter *s*.
4) Lipolysis is inversely proportional to systemic insulin (Jensen et al. 1989).
5) B cells, which produce insulin, increase proportionally to systemic glucose at low concentrations; at high concentrations, glucose is toxic, a phenomenon known as glucotoxicity (Efanova et al. 1998).
6) There exists glucose storage capacity, described by parameter K; excess glucose is assumed to be primarily stored in adipose tissue and skeletal muscle. Here, we introduce a modification, where carrying capacity K is not a fixed parameter but instead a variable, which can decrease over time to mimic, for instance, age-related loss of muscle mass. A functional form

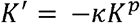

will achieve this property, with parameter kappa affecting the steepness of the decay, and parameter p will result in exponential decay if p=1, sublinear (slower that exponential) if p<1, and superlinear (faster than exponential) is p>1.

The modified system of equations will then become:

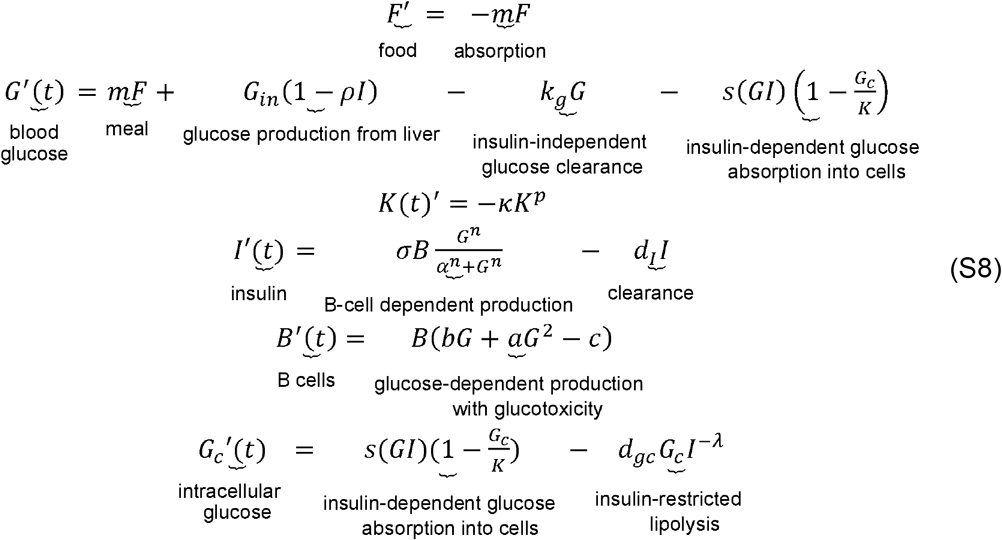

Parameters used for this updated model are summarized in Table S1.

**Table S1.** Parameter values used in System (S8).

| Parameter | Description | Value | Units | Refs |
| --- | --- | --- | --- | --- |
| $G_{in}$ | Glucose production from the liver (gluconeogenesis) | 0.25 | mM/min | Estimated |
| $\rho$ | Insulin-dependent reduction of gluconeogenesis | 0.008 | 1/min | Estimated |
| $k_g$ | Insulin-independent glucose clearance | 0.016 | 1/min | Estimated |
| $s$ | Rate of insulin-dependent glucose absorption into cells (insulin sensitivity) | 0.0008 | 1/uIU/L | (Kahn et al. 1993) |
| $\sigma$ | Rate of B-cell dependent insulin production | 0.06 | 1/min | (Topp et al. 2000) |
| $n$ | Parameter of steepness of the Hill function describing glucose-dependent insulin secretion by $\beta$ cells | 3.2 | n/a | (Alcazar & Buchwald 2019) |
| $\alpha$ | Glucose concentration for half-maximal insulin production by B cells | 8 | mM | (Alcazar & Buchwald 2019) |
| $d_I$ | Normal insulin clearance | 0.07 | 1/min | Estimated |
| $b$ | Rate constant for $\beta$ cell glucotoxicity | $5.833e-7$ | $1/\text{mM}^2/\text{min}$ | (Topp et al. 2000; Efanova et al. 1998) |
| $a$ | Rate constant for glucose-mediated $\beta$ cell expansion | $0.27e-4$ | $1/\text{mM}/\text{min}$ | (Topp et al. 2000; Efanova et al. |
|  |  |  |  | 1998) |
| c | B cell death rate | 1.667e-9 | 1/min | (Topp et al. 2000; Efanova et al. 1998) |
| d<sub>gc</sub> | Rate of insulin-restricted lipolysis | 0.035 | uIU/L/min | Estimated from (Jensen et al. 1989) |
| $\lambda$ | Scaling exponent that that controls the nonlinearity of the insulin-dependent lipolysis | 0.4 | n/a | Estimated from (Jensen et al. 1989) |
| $\kappa$ | Rate of glucose storage capacity decrease over time | 0.002 | 1/min | n/a |
| p | Scaling exponent that controls the nonlinearity of glucose storage capacity decrease over time | 0.94 | n/a | n/a |

Simulation of representative model dynamics over time with the introduced modification of the equation for age-related decline in the capacity to store glucose (which may arise, for instance, from age-related loss of muscle mass), is shown in Figure S3.

**Figure S3.**
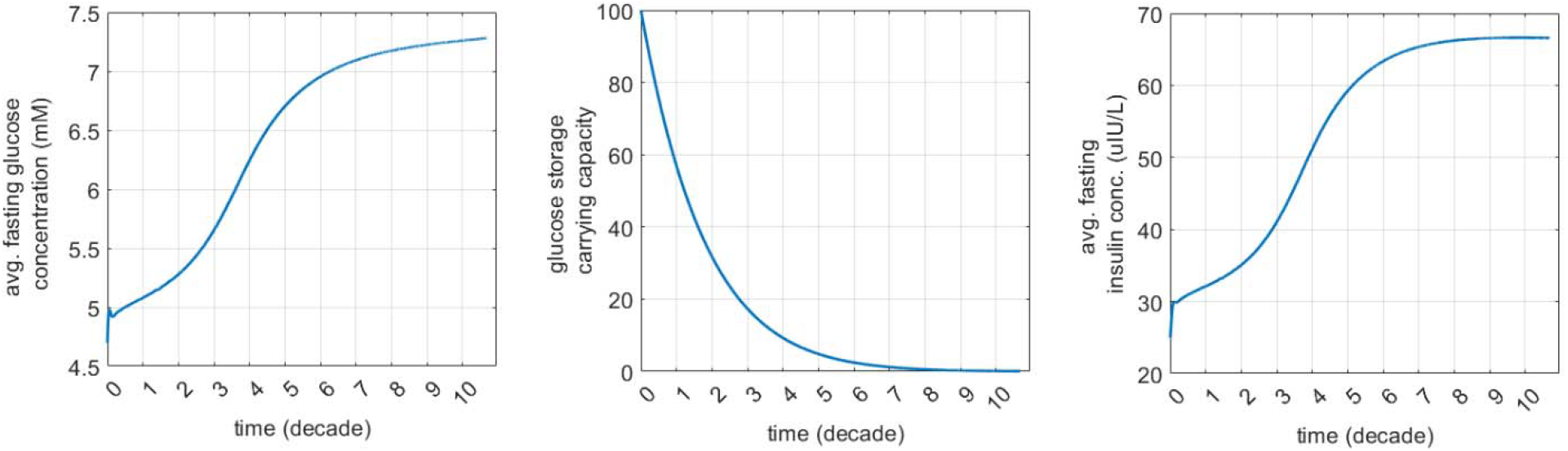
Representative dynamics of (a) average fasting glucose, (b) glucose carrying capacity that declines over time and (c) fasting insulin, capturing emergence of pre-diabetes and diabetes.

